# cyto: ultra high-throughput processing of 10x-flex single cell sequencing

**DOI:** 10.64898/2026.01.21.700936

**Authors:** Noam Teyssier, Alexander Dobin

## Abstract

Single-cell genomics is rapidly scaling toward billion-cell atlases, but computational analysis has become a critical bottleneck. Processing multiplexed datasets with existing tools requires substantial computational resources and runtime that become prohibitive at scale. Here we present `cyto`, an ultra highthroughput processor for 10x Genomics Flex single-cell sequencing optimized for production-scale analysis. `cyto` exploits the fixed sequence geometry of Flex libraries through direct k-mer lookup rather than alignment-based mapping, and introduces IBU (Indexed-Barcode-UMI), a compact binary format for efficient read processing. `cyto` further leverages BINSEQ, a binary sequencing format that enables highly parallel parsing and overcomes the single-threaded limitations of gzip compression. On a benchmark 320,000-cell multiplexed dataset, `cyto` completes processing in 13 minutes compared to `CellRanger`’s 3.7 hours, a 16.5-fold speedup, while requiring 2.4-fold less memory and performing 5.6-fold less disk I/O. The 31.7-fold reduction in CPU-hours represents true algorithmic efficiency rather than parallelization alone. Critically, `cyto` maintains 99.85% concordance with `CellRanger` outputs, with identical cell type clustering in dimensionality reduction analyses. These performance improvements enable costeffective processing on smaller cloud instances and make previously prohibitive experiments computationally feasible. `cyto` is production-ready, open-source software that provides the computational foundation for atlas-scale single-cell projects and genome-wide perturbation screens.

## 1 Introduction

Single-cell genomics has rapidly scaled from profiling thousands of cells to millions, with ambitious projects now targeting billion-cell atlases to map cellular diversity across tissues, development, and disease^1,2^. However, computational analysis has not kept pace with experimental throughput. Standard processing tools like `CellRanger`^3^, while robust, require substantial computational resources and runtime that become prohibitive at scale. For a typical 320,000-cell multiplexed dataset, `CellRanger` requires nearly 4 hours and 168 GB of peak memory, constraints that multiply across the hundreds or thousands of samples required for atlas-scale projects. When experimental protocols can generate data faster than it can be analyzed, computation becomes the rate-limiting step in discovery. This bottleneck translates directly to increased cloud computing costs, delayed insights, and reduced experimental iteration speed, ultimately limiting the scope of questions that can be addressed with single-cell approaches.

The emergence of highly multiplexed technologies like 10x Genomics Flex has exacerbated this computational challenge while simultaneously expanding experimental possibilities. Flex enables profiling of formalin-fixed paraffin-embedded (FFPE) samples with hundreds of probe sets simultaneously, democratizing single-cell analysis of archived clinical specimens and precious samples. Similarly, largescale CRISPR perturbation screens (Perturb-seq) now routinely assay thousands of genetic perturbations across millions of cells to map gene function and regulatory networks^4–6^. Despite the growing adoption of Flex technology, `CellRanger` remains the only available tool for demultiplexing and processing these multiplexed datasets, creating a dependency on software not optimized for production-scale throughput. As experimental costs per cell decrease and throughput increases, computational costs increasingly dominate project budgets. Cloud computing expenses scale linearly with runtime and memory usage, making the processing of large multiplexed datasets a significant operational burden for research institutions and production facilities. This computational bottleneck is further compounded by the lack of alternative tools, limiting researchers’ ability to optimize their processing pipelines for cost and speed. Addressing these constraints is essential not just for cost reduction, but for enabling the rapid experimental iteration required for large-scale functional genomics.

Here we present `cyto`, an ultra high -throughput processor for 10x Flex single-cell sequencing designed specifically for production-scale analysis. `cyto` achieves dramatic performance improvements through algorithmic innovations that exploit the fixed sequence geometry of Flex libraries. Rather than alignmentbased read mapping, `cyto` uses direct k-mer lookup against pre-computed hash tables, eliminating the computational overhead of traditional alignment algorithms. We introduce IBU (Indexed-Barcode-UMI), a compact binary format that enables efficient read tracking and processing between pipeline stages, replacing the inefficient text-based intermediate files common in existing workflows. `cyto` further leverages BINSEQ, a binary sequencing format that enables highly parallel sequence parsing, overcoming the single-threaded limitations of gzip compression^7^. The modular architecture allows independent scaling of pipeline components and straightforward customization for specialized experimental designs. On a benchmark 320,000-cell multiplexed FFPE dataset, `cyto` completes processing in 13 minutes compared to `CellRanger`’s 3.7 hours, a 16.5-fold speedup, while requiring 2.4-fold less memory and performing 5.6-fold less disk I/O. Critically, this performance gain comes without sacrificing accuracy: single-cell transcriptomes show 99.85% concordance with `CellRanger` outputs across all samples.

These performance improvements have immediate practical implications for large-scale single-cell projects. The 31.7-fold reduction in CPU-hours represents true algorithmic efficiency rather than simple parallelization, enabling cost-effective processing on smaller cloud instances and significantly reducing per-sample computational costs. For atlas-scale projects processing thousands of samples, these savings translate to substantial reductions in both infrastructure costs and time-to-insight. More importantly, `cyto`’s efficiency enables experimental designs that were previously computationally prohibitive. The modular architecture naturally extends to upcoming higher-plex Flex chemistries (364-plex in Flex-V2) where existing tools would face even more severe scaling challenges. For Perturb-seq experiments, `cyto` includes an efficient guide assignment algorithm (geomux) that scales linearly with dataset sparsity rather than with total cell and guide counts, enabling million-cell perturbation screens without requiring prefiltering or subsampling. By removing computational bottlenecks, `cyto` allows experimental throughput to drive project timelines rather than analysis capacity, fundamentally changing the economics and feasibility of large-scale functional genomics studies. `cyto` is implemented as an open-source commandline tool, providing the single-cell genomics community with a production-ready alternative for highthroughput Flex processing that matches the scale and speed of modern experimental platforms.

## 2 Results

We benchmark `cyto` against `CellRanger` (version 9.0.1) on the 320k scFFPE from 8 Human Tissues 320k, 16-Plex dataset. Both tools were run on identical hardware, on NVMe SSDs, using 128 threads.

### 2.1 Runtime and resource statistics

`cyto` completed processing in 13 minutes compared to CellRanger’s 3.7 hours, showing a 16.5-fold speedup (Figure 1a). This performance gain extended across all resource dimensions: `cyto` consumed 4.3 CPU-hours versus 138.5 CPU-hours (31.7x reduction, Figure 1c), required 70 GB peak memory versus 168 GB (2.4x reduction, Figure 1b), and performed 689 GB of disk I/O versus 3837 GB (5.6x reduction, Figure 1d).

**Figure 1:**
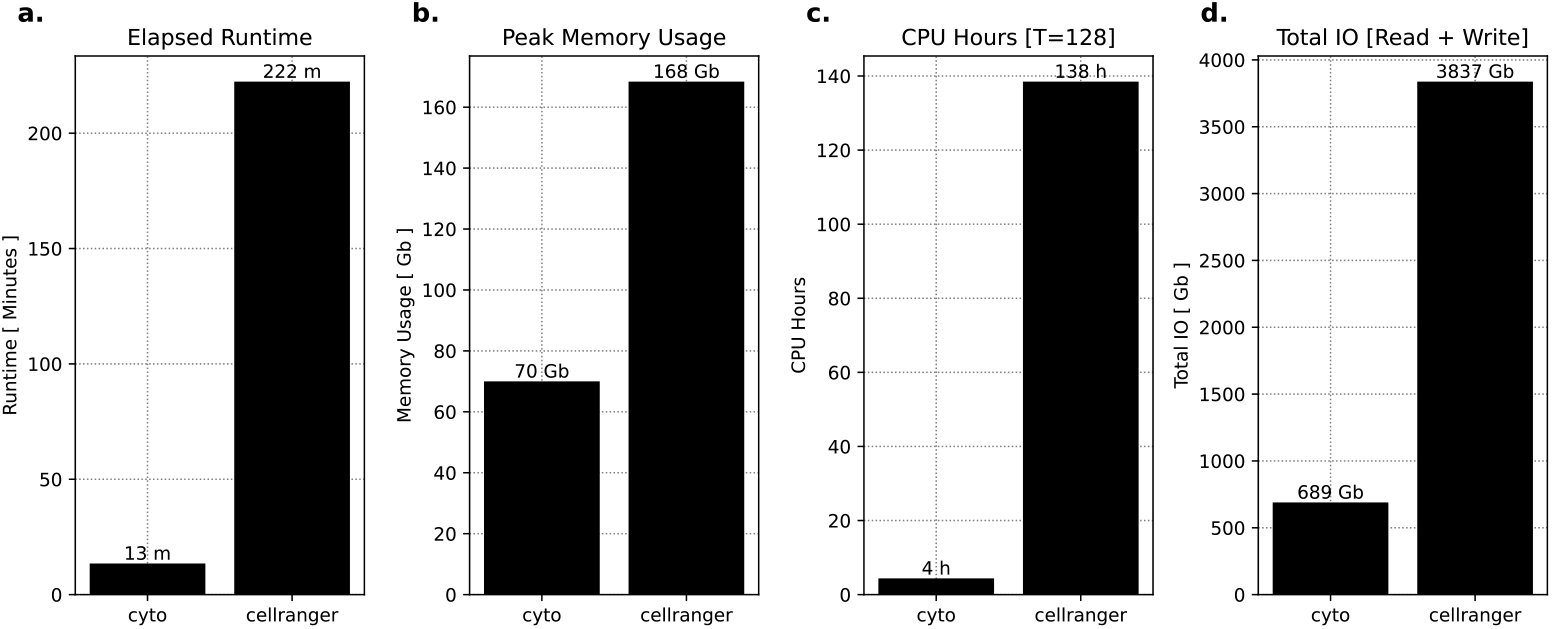
Runtime statistics for `cyto` and `CellRanger` on the 320k scFFPE from 8 Human Tissues 320k, 16-Plex dataset. (a) Elapsed time in minutes, `cyto` shows a 16.5x speedup over `CellRanger`. (b) Peak memory usage in GB, `cyto` shows a 2.4x reduction over `CellRanger`. (c) CPU hours, measured in hours, `cyto` shows a 29.7x efficiency over `CellRanger`. (d) Total IO measured in GB, `cyto` shows a 3.9x reduction in IO compared to `CellRanger`.

This dramatic reduction in CPU-hours indicates true algorithmic efficiency rather than simple parallelization, as `cyto` achieved higher throughput while using proportionally fewer computational resources. Peak memory reduction enables deployment on more cost-effective cloud instances, while the 5.6x decrease in disk I/O reduces both runtime on I/O constrained systems and storage infrastructure costs.

### 2.2 Modular pipeline architecture enables consistent performance

To assess the consistency and scalability of `cyto`’s architecture we examined per-module runtimes across the 16-Flex probe barcodes that were processed in parallel (Figure 2). Mapping required only 187 seconds of the total 13 minute runtime (23% of runtime) resulting in a 34M reads per second throughput. This throughput is achieved in part by leveraging the parallelization capabilities of BINSEQ, dramatically improving upon the single-threaded limitations of gzip FASTQ. Notably, this also improves upon CellRanger, which in order to achieve parallelization of FASTQ shards the input files into a high number of smaller LZ4 compressed chunks of FASTQ. That approach is significantly less efficient than `cyto`’s approach as it both contributes to very large I/O overhead, disk usage, and increases the runtime dramatically.

**Figure 2:**
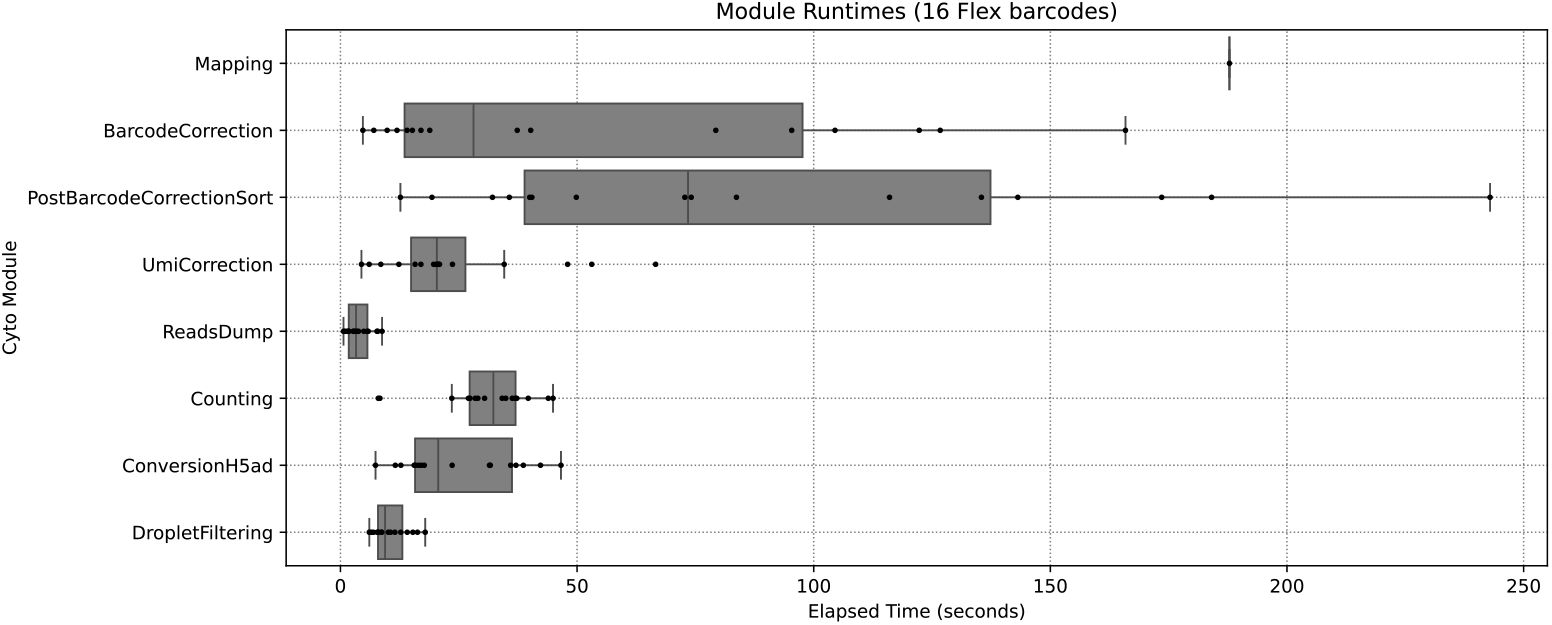
Individual module runtime statistics for `cyto`. Read mapping is done once for all flex barcodes and all downstream modules can be run in parallel across each flex barcode separately. This workflow can be parallelized by `cyto` but can also be handled explicitly by the user for improved performance and resource management via standard orchestration tools.

The remaining modules are each run in parallel for each of the barcodes separately (16 independent workflows). Some of the modules, such as sorting and UMI correction, can be run with additional threads, and `cyto`’s current implementation splits the allotted threads across the barcodes evenly without any work-stealing. The most expensive steps are Barcode Correction (i.e. whitelist comparison and 1-hamming distance correction) and the sort step that is required afterwards. The variance observed in these steps correlates strongly with the number of records observed in those individual barcodes, which is not guaranteed to be uniform across all barcodes.

Each of these steps does not benefit much from additional parallelization and as such can scale well to additional Flex barcodes (such as the 364 expected in Flex-V2).

### 2.3 Single-Cell transcriptome concordance

To validate that performance improvements did not compromise analytical accuracy, we compared `cyto` and CellRanger outputs across the 8 diverse scFFPE datasets spanning multiple tissue types (lung, breast, colorectal, glioblastoma, kidney, endometrium, skin melanoma, and lymph node). Cell-level per-gene expression values showed exceptionally high correlation across all samples (mean Spearman *ρ* = 0.998, Figure 3a) with cell counts ranging from 15k to 66k per sample.

**Figure 3:**
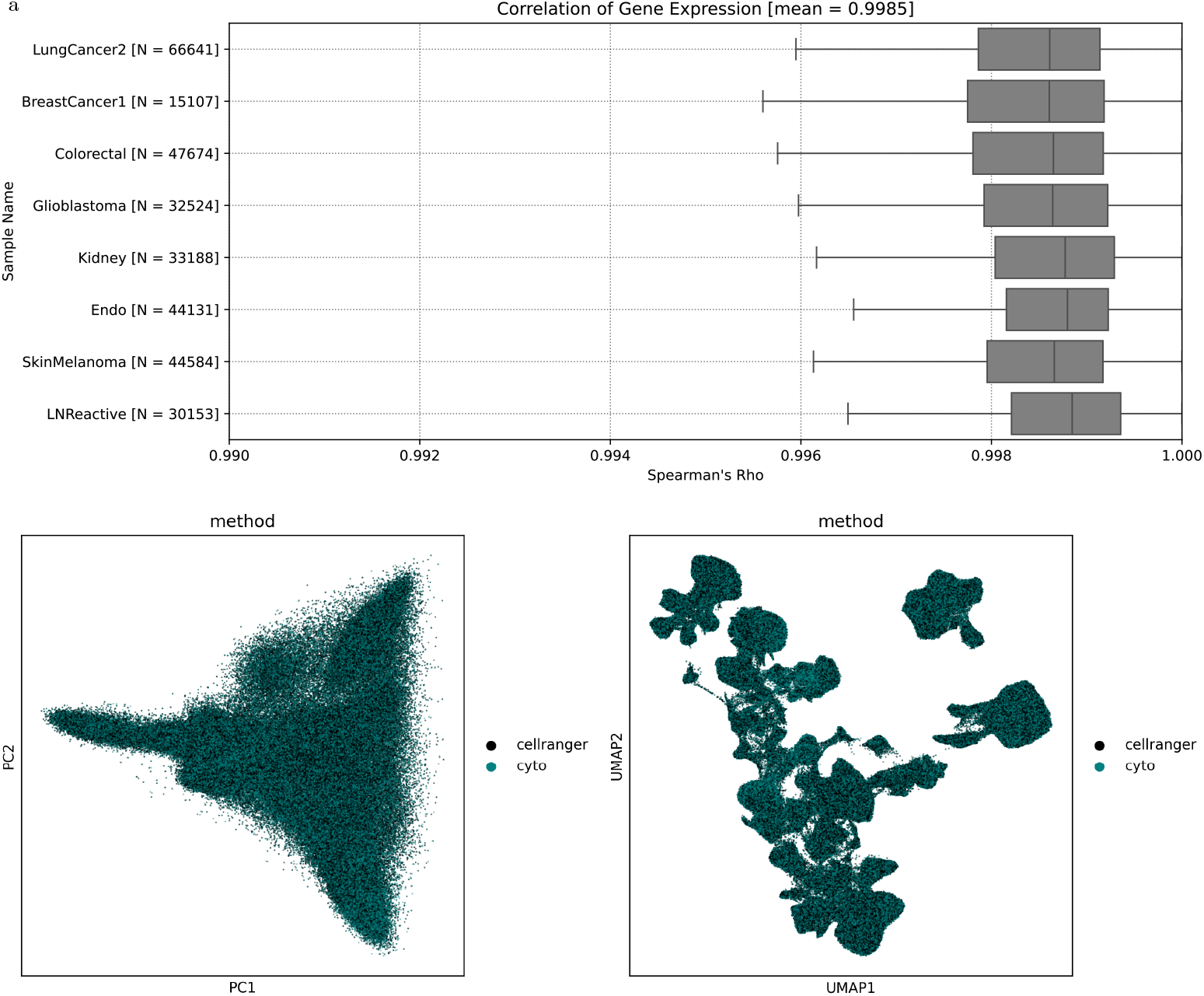
Comparison of output single-cell transcriptomes between `cyto` and `CellRanger`. (a) Spearman correlation of individual cell gene expressions show an average of 99.85% across all samples. (b) PCA and (c) UMAP plots show identical clustering of cells without any method-specific bias.

Dimensionality reduction via PCA (Figure 3b) and UMAP (Figure 3c) reveal complete overlap between methods in both projections with no systematic differences in cell type clustering or transcriptional structure. The near-perfect correlation and identical low-dimensional structure demonstrate that `cyto`’s algorithmic optimizations preserve biological signal while dramatically reducing computational burden.

## 3 Methods

### 3.2 Analysis Workflow

`cyto` is implemented in rust and is composed of modular components that can be extended, customized, or replaced. This design allows for flexibility of analysis, extensibility, and potential for optimizations on different hardware architectures.

`cyto` exposes all components as independent command-line tools which can be run independently. However, it also provides workflow commands which orchestrate the execution of commonly used pipelines for analysis. `cyto` also introduces a simple and efficient binary file, IBU (Indexed-Barcode-UMI), which is used for tracking, storing, and transmitting information about reads between the components in a compact binary format.

The major components of `cyto` include: Read mapping, IBU sorting, Barcode Correction, UMI Correction, Transcript Counting, Droplet Filtering, and Guide-Calling.

#### 3.1.1 Read Mapping

`cyto` performs read mapping through direct k-mer lookup rather than alignment-based sequence matching. This approach exploits the fixed sequence geometry of 10X Flex libraries, where probe sequences and demultiplexing barcodes appear at predetermined positions within sequencing reads. By extracting subsequences at known offsets, `cyto` achieves efficient mapping without computationally expensive methods used in traditional alignment-based approaches.

`cyto` implements two mapping modes to handle different probe architectures: *gex* for transcript-targeting probes and *crispr* for guide RNA-targeting constructs.

In *gex* mode, the R2 read begins with the transcript-targeting probe sequence, followed by a demultiplexing barcode at a fixed downstream position. In *crispr* mode, the R2 read begins with the demultiplexing barcode, followed by an anchor sequence and protospacer at fixed downstream positions. The length of constant regions (adapters, spacers) between these elements is configurable at runtime to accommodate different experimental designs.

### Index Construction and Lookup Strategy

Prior to processing reads, `cyto` constructs hash-based lookup tables for all library elements in both the probe library (transcript probes or protospacers) and demultiplexing barcode library. Each unique sequence is mapped to a unique integer index representing its position in the library.

To enable mismatch-tolerant mapping, `cyto` generates a secondary mismatch table containing all 1-Hamming distance variants of each library sequence. Critically, variants that are 1-Hamming distance from multiple library elements are excluded from this table to prevent ambiguous assignments. For example, probe sequences ACGT and ACGC differ by only one base so neither ACGG nor other ambiguous 1-mismatch variants are included in the mismatch table. This ensures deterministic mapping: each query sequence maps to at most one library element.

This pre-computed mismatch table enables *O*(1) lookup operations for sequences with single base substitutions, avoiding expensive dynamic alignment or edit distance calculations. However, this approach does not capture insertions or deletions within the probe sequence itself. Mismatch tolerance can be disabled at runtime if deterministic exact matching is required.

### Read Mapping Procedure

For each read pair, `cyto` extracts the probe and demultiplexing barcode sequences at their expected positions based on the configured geometry. Each extracted sequence is queried against its corresponding exact-match lookup table via hash lookup. If an exact match is found, the read is assigned to that library element’s index.

If no exact match is found and mismatch tolerance is enabled, `cyto` queries the 1-mismatch table. Sequences absent from both tables are considered unmapped.

To account for insertions or deletions in constant regions (adapter sequences, spacers) that shift the expected positions of probe and barcode sequences, `cyto` implements an optional remapping procedure. When enabled, unmapped sequences are re-extracted at ±1bp offsets from the configured positions and queried against the lookup tables. This handles cases where library preparation artifacts introduce small indels outside the probe sequence itself.

Successfully mapped reads are written to sample-specific output files organized by demultiplexing barcode. Each record stores the cell barcode, UMI, and the integer index corresponding to the mapped probe in a compressed binary format (IBU format, see Supplementary Methods).

### Mapping Gene Expression Probes

The gene expression probes are quite error prone and are mapped in a different manner than the CRISPR or demultiplexing probes. `cyto` uses a similar algorithm to the approach taken by `CellRanger`.

The major difference is that gene expression probe libraries are generated from the first and last half of the expected probe sequence separately. Each incoming subsequence is then split in half and queried against both the exact-match and 1-mismatch lookup tables for their respective halves. If both halves correctly map (or are one-mismatch away), their indices are compared and if they match (both indices are equal), the read is considered mapped to that index and written to its corresponding IBU. In any other case, the read is considered unmapped. This approach will additionally capture reads with less than three mismatches (though not all depending on the mismatches placement).

Notably, `cyto` does not track half-mapped reads in the same way that `CellRanger` does, as it only considers reads mapped or unmapped. However the alignment statistics are tracked and reported in the mapping step which does track these statistics.

#### 3.1.2 IBU And IBU Sorting

The IBU format is a compressed binary format used by `cyto` to track cell-barcode, UMI, and an integer index corresponding to a library element for a mapped read. The format is designed to be very efficiently serializable and deserializable, store and compress well, and is optimized for simple operations over large collections. Notably the individual records of the format can fit on the stack easily and efficiently. It is inspired by the BUS format originally developed by the Melsted et al., but simplified even further to improve IO performance^8^.

The IBU format begins with a header which annotates the length of each cell-barcode and the length of each UMI. Then each record is exactly 24 bytes long, consisting of 3 64-bit unsigned integers. The first integer is the two-bit encoded barcode sequence. The second integer is the two-bit encoded UMI sequence. The third integer is the library index that read mapped to.

IBU sorting is a crucial step in the processing pipeline that occurs a few times during processing. IBUs are generated per probe-element (a maximum of 16 for Flex-V1) and as such are expected to be roughly 1/16th the total number of reads. Because these are fixed binary records, their size is deterministic (exactly 24 bytes per record), so for a 1B read set each of the 16 IBUs will be approximately 1.4Gb.

Oftentimes each of these files can be easily loaded into memory, allowing for efficient sorting and processing with in-memory algorithms. However, for larger datasets, or when processing large numbers of samples, it is often necessary to use out-of-memory sorting algorithms to reduce memory usage and increase the amount of concurrent operations available.

`cyto` has built-in support for out-of-memory sorting of IBU files using an external merge sort algorithm. This algorithm is built on the efficient serialization and deserialization of IBUs and allows for limiting the maximum memory per process at the cost of increased disk I/O. It also makes use of parallel processing to increase throughput.

Notably, IBU files are uncompressed binary files by default, but can be additionally compressed with standard compression algorithms such as gzip, bzip2, or zstd. Practically we find that these files have a compression ratio of approximately 40-45% on average.

#### 3.1.3 Barcode Correction

`cyto` employs a barcode-correction algorithm that can correct single-base errors in cell barcodes. It accepts a whitelist of valid barcodes and generates a disambiguated mismatch table from it in the same manner as in the mapping algorithm.

Then for each record we query this table with the barcode. For barcodes that are in the whitelist, we write the IBU to the output file. For barcodes that are one base away from a valid barcode, we write the corrected barcode to the output file. For barcodes that map to an ambiguous barcode (i.e. 1-hamming distance from two in the whitelist), we store the record in an auxiliary collection.

Once through the entire IBU, we reprocess the auxiliary collection and if only one of all ambiguous parent sequences was observed (all other ambiguous parents have zero occurrents) we can correct the barcode to that sequence. This allows rescuing of records with ambiguous mismatches where only one parent sequence was actually present. It accounts for a very small fraction of total records but can increase the global UMI abundance across valid barcodes. This secondary pass is optional and can be disabled at runtime.

#### 3.1.4 UMI Correction

UMI correction is a process that aims to correct potential mismatches in UMI sequences. This problem is more challenging than barcode correction because there is no whitelist of valid UMIs to correct to. As such, the correction process must rely on other UMIs in the same barcode that target the same index. If there are multiple UMIs in the same barcode that target the same index and those UMIs are within 1-hamming distance of each other, we can correct lower-abundance UMIs to the most abundant one.

Our implementation of UMI correction is based on finding connected components where each component represents a group of UMIs that are within 1-hamming distance of each other and belong to the same barcode and index. We first construct a graph for each barcode where each node represents a UMI sequence and an edge between two nodes indicates that the UMIs are within 1-hamming distance of each other. We then apply a depth-first search to find the connected components in the graph. For each component, we identify the most abundant UMI sequence and correct all other UMIs in the component to this sequence.

This algorithm is run and parallelized over each barcode independently. For ideal memory throughput the input is expected to be sorted by barcode before processing.

#### 3.1.5 UMI Counting and Deduplication

The final step of the sequencing pipeline is to iterate over all the sequencing reads in the IBUs, deduplicate UMIs, and then count the number of UMI-Index records per barcode. This is done in a cache-efficient manner by first sorting the IBUs by barcode and UMI and performing a one-pass counting algorithm that maximizes the number of hash lookups per barcode, UMI, and index.

The IBU records are first sorted lexicographically by barcode, then UMI, then index. This ensures that all records belonging to the same barcode-UMI combination are contiguous in the stream, enabling singlepass processing. We initialize a nested hash table that maps each barcode to an inner hash table, which in turn maps each index to its count. This structure stores the final deduplicated counts. During iteration, we maintain state tracking the current index being observed and its abundance (number of observations), the maximum observed abundance for the current UMI group and its corresponding index, and whether multiple indices share the maximum abundance.

The algorithm begins by selecting the first record in the IBU stream as the initial reference. For each subsequent record, we perform a series of comparisons to determine the appropriate action. When encountering a new barcode, we finalize the previous UMI group by comparing the current index abundance against the maximum observed for that UMI. If a single index has the highest abundance with no ties, we record one count for that barcode-index pair. The tracking state is then reset to begin processing the new barcode. When the barcode matches but a new UMI is encountered, we apply the same finalization process before resetting state to track the new UMI. When both barcode and UMI match but the index differs, we compare the previous index’s abundance against the current maximum before beginning to track the new index with an initial abundance of one. If all three fields match, we increment the abundance counter for the current index. After each comparison, the reference record is updated to the current record.

When multiple indices within a UMI group have equal maximum abundance, the UMI is discarded and no count is recorded for that barcode-UMI combination. This conservative approach avoids ambiguous index assignments that could arise from sequencing errors or PCR artifacts. After processing all records, we perform a final comparison and conditional count insertion to handle the last UMI group in the stream. The output is a sparse matrix representation of barcode-index counts, where each count represents a unique UMI that unambiguously mapped to that index.

The IBU count submodule is also used to aggregate the UMI counts of probes that share a gene group. The algorithm above is used to calculate the UMI count for each barcode-index pair. Then for each barcode-index pair we sum the UMI counts for all indices that belong to a gene group. We then report the UMI counts for each barcode-group pair.

#### 3.1.6 Cell Filtering

To account for empty droplets recovered by the sequencing protocol we include a similar implementation of the cell filtering algorithm employed by `CellRanger`. This algorithm is based on the `EmptyDrops` algorithm originally proposed by Lun et al. but with a few notable differences^9^. This is reimplemented and distributed as a standalone python application, `cell-filter`, which is used internally by `cyto` but can be run generically on other single-cell datasets. This algorithm is run by default in the workflow but can be disabled if alternative cell-calling methods are preferred.

A key differences in the `cell-filter` algorithm is how the thresholds for the putative set of cell-barcodes are selected. `EmptyDrops` calculates a knee point through a minimization algorithm, while the approach taken by `CellRanger` and `cell-filter` uses a ratio of the percentiles of the UMI counts. Another key difference is the distribution used for testing the null hypothesis. `EmptyDrops` uses a Dirichlet-Multinomial distribution for evaluating the transcriptome of each putative cell barcode, while by default `CellRanger` uses a Multinomial distribution. By default `cell-filter` uses a Multinomial to reproduce the approach taken by `CellRanger`, but provides both Dirichlet-Multinomial and Multinomial implementations for generic use.

#### 3.1.7 Guide Assignment

`cyto` supports perturb-seq experiments and creates cell-x-guide matrices. An important use case for these matrices is to assign cells to guides for downstream analysis. For many perturb-seq contexts the number of perturbations per guide is not fixed and can depend heavily on the experimental design. Many guide assignment algorithms focus on assignment of a single-perturbation per cell which is not applicable in multi-guide perturbations common in perturb-seq experiments. For CRISPR data this algorithm is run by default as it is very efficient and does not add significant computational overhead - however it can be disabled if a different algorithm is desired.

`cyto` makes use of the `geomux` algorithm for guide-assignment^10^. Briefly, a background distribution for each of the guides is calculated across the dataset. Then for each cell-guide pair we perform a hypergeometric test given the total UMIs of that cell, the background distribution of that guide, and the observed UMIs for the cell-guide pair. We then perform multiple-hypothesis correction using the Benjamini-Hochberg procedure. Finally a dynamic algorithm is used to further distinguish background from signal where for a given cell the log-odds ratio is calculated between the most significant non-significant guide and the most insignificant significant guide. If this log-odds ratio does not meet a given threshold then the significant status of the guide is dropped and the algorithm is repeated until the log-odds ratio threshold is met. This results in a set of assignments for each cell which can be filtered downstream to the expected multiplicity of infection and expected guide groups.

Notably, because the hypergeometric test needs only be performed on non-zero elements this algorithm has both computational and memory complexity of *O*(*n*) where *n* is the number of non-zero elements. In practice, because cell-x-guide matrices in practice are hyper sparse, this algorithm can approach 1000x throughput to existing algorithms which generally scale *O*(*NM*) where *N* is the number of cells and *M* is the number of guides.

A benefit of this algorithms low resource requirements is that it does not need to be run on a subset of cells (i.e. filtered cells from gene expression data). When running a perturb-seq workflow with `CellRanger` the Gaussian-Mixture-Model approach must be run on the filtered cells because performing the algorithm on unfiltered cells is computationally unfeasible and may introduce errors. Using the `geomux` algorithm we do not need to borrow information between modalities and can efficiently assign guides to cells without filtering *a priori*.

### 3.2 Benchmarking Details

To compare `cyto` against the existing standard, `CellRanger` we ran a comparison between the two programs on the 320*k scFFPE From* 8 *Human Tissues* 320*k*, 16-*Plex* dataset from 10X Genomics. The four input FASTQ pairs provided by 10X Genomics total 433GB of sequences (average 33GB for R1, 74GB for R2). Their corresponding BINSEQ (vbq) equivalents are 271GB (average 67GB each). The collection includes a total of 7.3 billion records.

We make our comparison against `CellRanger` version 9.0.1 on a generic linux server with 128 cores, 1T of total RAM, and SSD storage. Our `CellRanger` config sets `create_bam,false` and `no-secondary,true` to minimize resource usage and make a closer comparison. Both programs were run with a cold cache to simulate true production environments.

Runtime statistics were gathered for `cyto` using `/usr/bin/time -v` which tracks kernel-level resource usage including resident memory, CPU time, and block-level I/O operations. `CellRanger` resource statistics were extracted from its internal performance logs (`_perf.json`), using the `SC_MULTI_CS` stage metrics for memory (`highmem.rss`), CPU hours (`core_hours`) and I/O operations (`total_blocks`). Block-level I/O was converted to bytes using the standard 512-byte block size.

## 4 Discussion

`cyto` achieves a 16.5-fold reduction in runtime and 31.7-fold reduction in CPU-hours compared to `CellRanger`, representing true algorithmic efficiency rather than simple parallelization. Critically, this performance gain preserves analytical accuracy: single-cell transcriptomes show 99.85% concordance with `CellRanger` outputs, with identical cell type clustering in dimensionality reduction analyses. These results demonstrate that domain-specific optimization can overcome computational bottlenecks in single-cell genomics without sacrificing biological accuracy.

`cyto`’s performance stems from exploiting the fixed sequence geometry of Flex libraries. Direct k-mer lookup eliminates alignment overhead when probe and barcode positions are predetermined. The IBU binary format represents each read in exactly 24 bytes versus hundreds required by text-based formats, reducing both disk I/O (5.6-fold) and memory overhead (2.4-fold). BINSEQ enables efficient parallel parsing, avoiding `CellRanger`’s approach of sharding FASTQ into numerous chunks that increases I/O dramatically. With mapping consuming only 23% of runtime, the modular architecture scales naturally to higher-plex chemistries like Flex-V2′s 364-plex format.

These improvements enable processing on smaller cloud instances and reduce per-sample computational costs substantially. For atlas-scale projects processing thousands of samples, this translates to significant infrastructure savings and faster time-to-insight. More importantly, `cyto` enables previously prohibitive experiments. The geomux guide assignment algorithm scales linearly with dataset sparsity rather than total dimensions, enabling million-cell perturbation screens without pre-filtering that could introduce bias. By removing computational bottlenecks, `cyto` allows experimental throughput rather than analysis capacity to drive project timelines.

`cyto` is optimized for fixed-geometry protocols and is not a general-purpose aligner. The k-mer approach requires known sequence structures, making it unsuitable for splice-aware alignment, transcript discovery, or variable read architectures - applications where tools like STAR, kallisto|bustools, and Alevin-Fry remain appropriate^11,12,13^. The 1-hamming distance tolerance does not capture insertions or deletions within probes themselves, though optional ±1bp offset remapping handles indels in constant regions. These design choices reflect `cyto`’s focus on production-scale processing of well-characterized libraries.

The modular architecture enables continued optimization and extension. The IBU format could adapt to other multiplexed technologies with fixed geometries. Specific modules could leverage GPU acceleration or SIMD optimizations. As single-cell technologies scale toward million-sample studies and billion-cell datasets, tools prioritizing algorithmic efficiency will become essential infrastructure. The open-source implementation enables community contributions and domain-specific customizations.

`cyto` addresses a critical computational bottleneck in single-cell genomics through algorithmic innovations designed for modern multiplexed technologies. By achieving order-of-magnitude performance improvements without compromising accuracy, `cyto` removes barriers limiting the scope and pace of large-scale functional genomics. `cyto` is freely available as open-source software, providing the computational foundation needed to match analytical throughput with experimental capacity as the field moves toward atlas-scale projects and genome-wide perturbation screens.

## 5 Software Availability

All software components are free and open-source. `cyto` is available on GitHub as https://github.com/arcinstitute/cyto. `pycyto` provides related utilities, such as aggregation and format conversions, and is available on GitHub as https://github.com/arcinstitute/pycyto. `geomux` is available on GitHub as https://github.com/noamteyssier/geomux and includes both the `geomux` algorithm as well as the Gaussian-Mixture model (GMM) implementation. `cell-filter` is available on GitHub as https://github.com/arcinstitute/cell-filter. The IBU format description and rust library are available on GitHub as https://github.com/noamteyssier/ibu.

All rust components are distributed via `crates.io` and python components are available on `PyPI` and packaged via `uv`.

## 6 Author Contributions

N.T. designed and implemented `cyto`, performed benchmarking analyses, and wrote the manuscript. A.D. supervised the project.

## 8 Data Availability

The benchmark dataset (320k scFFPE from 8 Human Tissues, 16-Plex) is publicly available from 10X genomics (https://www.10xgenomics.com/datasets/320k_scFFPE_16-plex_GEM-X_FLEX).

## 9 Acknowledgements

We thank the Arc Institute for computational resources and support.

